# A strain-specific multiplex RT-PCR for Australian rabbit haemorrhagic disease viruses uncovers a new recombinant virus variant in rabbits and hares

**DOI:** 10.1101/173930

**Authors:** Robyn N Hall, Jackie E. Mahar, Andrew J. Read, R Mourant, M Piper, N Huang, Tanja Strive

**Affiliations:** CSIRO, Acton, ACT, 2601, Australia; Invasive Animals Cooperative Research Centre, University of Canberra, Bruce, ACT, 2601, Australia; The University of Sydney, Sydney, NSW, 2006, Australia; Elizabeth Macarthur Agricultural Institute, Menangle, NSW, 2568, Australia

**Keywords:** Caliciviridae, Lagovirus, Polymerase Chain Reaction, Hemorrhagic Disease Virus, Rabbit, Recombination, Genetic

## Abstract

*Rabbit haemorrhagic disease virus* (RHDV, or GI.1), is a calicivirus in the genus *Lagovirus* that has been widely utilised in Australia as a biological control agent for the management of overabundant wild European rabbit (*Oryctolagus cuniculus*) populations since 1996. Recently, two exotic incursions of pathogenic lagoviruses have been reported in Australia; GI.1a-Aus, previously called RHDVa-Aus, is a GI.1a virus detected in January 2014, and the novel lagovirus GI.2 (previously known as RHDV2). Furthermore, an additional GI.1a strain, GI.1a-K5 (also known as 08Q712), was released nationwide in March 2017 as a supplementary tool for wild rabbit management. To discriminate between these lagoviruses, a highly sensitive strain-specific multiplex RT-PCR assay was developed, which allows fast, cost-effective, and sensitive detection of the four pathogenic lagoviruses currently known to be circulating in Australia. In addition, we developed a universal qRT-PCR assay to be used in conjunction with the multiplex assay that broadly detects all four viruses and facilitates quantification of viral RNA load in samples. These assays enable rapid detection, identification, and quantification of pathogenic lagoviruses in the Australian context. Using these assays, a novel recombinant lagovirus was detected in rabbit tissues samples, which contained the non-structural genes of GI.1a-Aus and the structural genes of GI.2. This variant was also recovered from the liver of a European brown hare (*Lepus europaeus*). The impact of this novel recombinant on Australian wild lagomorph populations and its competitiveness in relation to circulating field strains, particularly GI.2, requires further studies.

## Introduction

Rabbit haemorrhagic disease (RHD) is caused by pathogenic rabbit caliciviruses belonging to the genus *Lagovirus*. RHD affects European rabbits of the genus *Oryctolagus* and is characterised by a necrotising hepatitis leading to multi-organ failure and death, frequently within 48-72 hours post-infection (Abrantes *et al*., 2012). RHD was first reported in China in 1984 and was later recognised as being caused by the rabbit calicivirus GI.1 (previously referred to as *Rabbit haemorrhagic disease virus* or RHDV) (Liu *et al*., 1984, Ohlinger *et al*., 1990, Le Pendu *et al*., 2017). Subsequently, in the late 1990s, GI.1 variants were reported that had unique reactivity profiles with monoclonal antibodies and were genetically divergent from previously sequenced GI.1 viruses (Capucci *et al*., 1998, Schirrmeier *et al*., 1999, Le Gall-Recule *et al*., 2003). These antigenically distinct variants were designated GI.1a (previously referred to as subtype RHDVa or genogroup G6), while classical GI.1 viruses were sub-classified into GI.1b, GI.1c, and GI.1d variants based on phylogenetic analyses (previously called genogroups G1-G5) (Capucci *et al*., 1998, Le Gall-Recule *et al*., 2003, Lavazza and Capucci, 2016). In 2010, a novel lagovirus, GI.2 (previously called RHDV2), was reported in Europe (Le Gall-Recule *et al*., 2011a, Dalton *et al*., 2012). Although this virus also caused RHD, it was both antigenically and genetically divergent from both GI.1 and GI.1a viruses (Dalton *et al*., 2012, Le Gall-Recule *et al*., 2013, Lavazza and Capucci, 2016). GI.2 is able to cause disease in rabbits vaccinated against GI.1 and also affects young rabbit kittens, which are normally highly resistant to RHD caused by GI.1 or GI.1a (Le Gall-Recule *et al*., 2013).

In addition to the pathogenic lagoviruses GI.1, GI.1a, and GI.2, several non-pathogenic rabbit caliciviruses (RCVs) have been described from various geographical localities, including Italy (unclassified), Ireland and France (GI.3), and Australia (GI.4) (Capucci *et al*., 1996, Forrester *et al*., 2007, Strive *et al*., 2009, Le Gall-Recule *et al*., 2011b, Le Pendu *et al*., 2017). RCVs mainly cause a benign subclinical infection of the small intestine, in contrast to the pathogenic lagoviruses, which are generally hepatotropic (Capucci *et al*., 1996, Hoehn *et al*., 2013).

All lagoviruses have a single-stranded, positive-sense RNA genome of approximately 7.5 kb, and share a similar genome structure comprising two open reading frames (ORFs). ORF1 encodes a single polyprotein, which is cleaved into seven non-structural proteins and the major capsid protein VP60, while ORF2 encodes a minor structural protein, VP10 (Figure 1). The genome is polyadenylated at the 3’ end, and linked to a viral protein, VPg, at the 5’ end (Meyers *et al*., 1991). Additionally, these viruses produce a subgenomic RNA of approximately 2.2 kb that is also VPg-linked and polyadenylated (Meyers *et al*., 1991). This subgenomic RNA encodes both structural genes and is collinear with the 3’ end of the genomic RNA (Figure 1). This facilitates homologous recombination between the genomic and subgenomic RNAs, which frequently occurs at the junction between the non-structural and structural genes (i.e. at the RNA-dependent RNA polymerase (RdRp)-VP60 junction) (Meyers *et al*., 1991). Recombination appears to play an important role in generating genetic diversity in viruses of the *Caliciviridae* family, including the lagoviruses (Bull *et al*., 2007, Lopes *et al*., 2015). GI.1 recombinants have previously been reported (Forrester *et al*., 2008), and two GI.2 recombinants have been identified to date, a GI.1bP/GI.2 and GI.4P/GI.2 virus (Lopes *et al*., 2015), where the P denotes the RdRp genotype.

**Figure 1:**
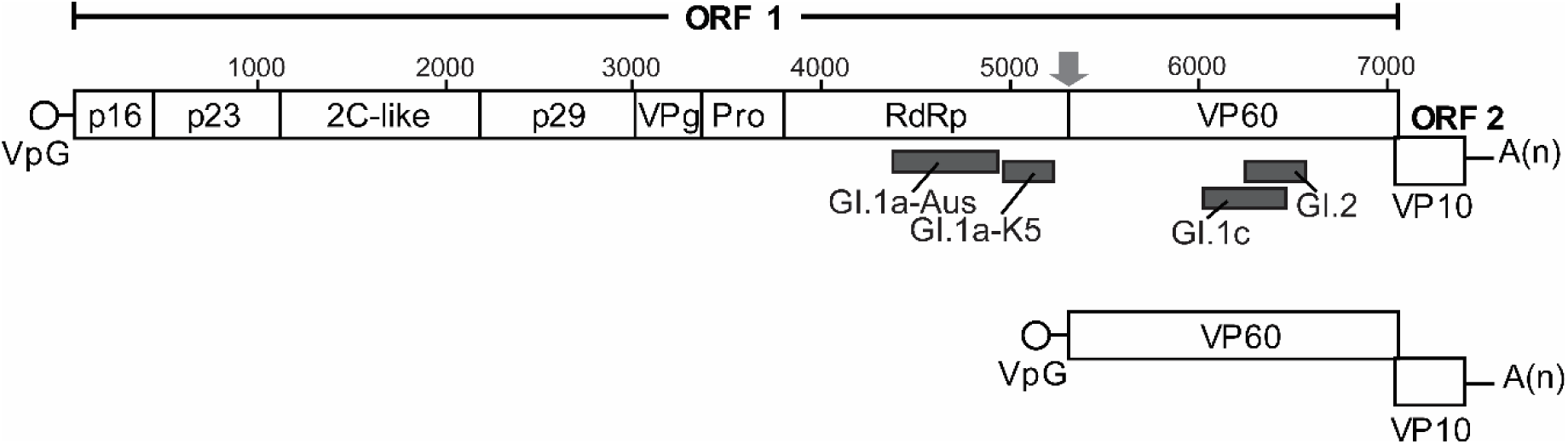
Genomic organisation of lagoviruses and location of multiplex RT-PCR amplicons. Lagoviruses have a single-stranded positive-sense RNA genome (top) approximately 7.5 kb in length comprising two open reading frames (ORFs; open boxes, labelled in bold). The ORF1 polyprotein is cleaved into seven non-structural proteins: p16, p23, 2C-like protein, p29, the viral genome-linked protein (VPg), the viral protease (Pro), the RNA-dependent RNA polymerase (RdRp), and the major capsid protein VP60. ORF2 encodes a minor structural protein, VP10. A subgenomic RNA (bottom) is also produced that is approximately 2.2 kb in length and encodes both structural proteins, VP60 and VP10. Both the genomic and subgenomic RNAs are polyadenylated at the 3’ end, and linked to VPg at the 5’ end. Recombination is frequently observed at the RdRp-VP60 junction (grey arrow). The regions amplified in the multiplex RT-PCR assay are shown (dark grey boxes) for GI.1a-Aus, GI.1a-K5, GI.1c, and GI.2 lagoviruses.

In Australia, GI.1 viruses are widely utilised as biological control agents for the management of wild European rabbits, which are a major invasive agricultural and environmental pest (Cooke and Fenner, 2002). A Czechoslovakian GI.1c strain (CAPM V-351) was first released for biocontrol purposes in the mid-1990s and this strain has been regularly re-released across Australia since this time (Cooke and Fenner, 2002, Kovaliski *et al*., 2014). Until 2014, the only lagoviruses known to be circulating in Australia were the benign *Rabbit calicivirus Australia*-*1* (RCV-A1; GI.4) and GI.1c field strains derived from the originally released GI.1c CAPM V-351 (Kovaliski *et al*., 2014, Eden *et al*., 2015, Mahar *et al*., 2016). However, in January 2014 an exotic GI.1a virus, GI.1a-Aus, was detected from an RHD outbreak in domestic rabbits in northern Sydney, New South Wales (NSW) (OIE, 2014). This virus subsequently caused multiple RHD outbreaks in NSW and the Australian Capital Territory (ACT) in both domestic and wild rabbits (RHDVa paper, in review). Full genome sequencing of GI.1a-Aus indicated that this was a recombinant virus, with a capsid gene most closely related to a 2010 GI.1a strain from China and non-structural genes similar to GI.4-like viruses (RHDVa paper, in review). Subsequently in May 2015, a second lagovirus incursion, GI.2, was detected in the ACT. This was also a recombinant virus (GI.2 capsid, GI.1b non-structural genes) that is closely related to viruses that circluated in Portugal and the Azores in 2014 (Hall *et al*., 2015). In addition, a Korean GI.1a strain, GI.1a-K5 (also known as 08Q712), was released nationwide in March 2017, to improve rabbit biocontrol program (Oem *et al*., 2009, Cox *et al*., 2013, OIE, 2017).

With the growing repertoire of lagoviruses now present in Australia, improved diagnostic tools are needed to enable rapid discrimination between the different viruses causing RHD and sudden death in both wild and domestic rabbits. Monitoring the spread and interactions of these viruses will help to maximise the effectiveness of wild rabbit control programs, and to guide the implementation of control measures for domestic rabbits. Here we report a sensitive and specific multiplex RT-PCR assay for the discrimination of the pathogenic lagoviruses circulating in Australia, namely classical GI.1c viruses, GI.1a-Aus, GI.1a-K5, and GI.2. Using this assay, we detected a new recombinant variant that has arisen from recombination between GI.1a-Aus and GI.2. Additionally, we describe a SYBR-green based qRT-PCR for the generic detection of all rabbit lagoviruses (GI.1c, GI.1a, GI.2, GI.4) circulating in Australia in a single PCR reaction. This assay facilitates quantification of viral RNA load in diagnostic samples from deceased animals, which is useful when interpreting equivocal results, since rabbits succumbing to fulminant RHD invariably have high viral loads in tissues (Elsworth *et al*., 2014), while low viral loads are likely to reflect sample contamination or residual vaccine virus. Taken together, the two assays allow robust and cost-effective sample analysis with a rapid turnaround time.

## Materials and Methods

### Virus samples

For multiplex RT-PCR assay development and validation, we used liver samples that had previously tested positive for different lagoviruses at the Elizabeth Macarthur Agricultural Institute (EMAI), Menangle, NSW. These samples had originally been collected from rabbits suspected to have died from RHD. Initial virus typing at EMAI was conducted using individual qRT-PCR assays specific for GI.1, GI.2, and GI.1a-Aus (Gall *et al*., 2007, Lavazza and Capucci, 2016) (RHDVa paper, in review). GI.1a-K5 was obtained from EMAI from stocks of the virus approved by Australian authorities for nationwide release. Liver samples and GI.1a-K5 stock virus were shipped to the Commonwealth Scientific and Industrial Research Organisation (CSIRO) on ice and stored at -20 °C on arrival.

For validation of the lagovirus qRT-PCR assay, in addition to the known lagovirus-positive samples from EMAI, liver or bone marrow samples from wild or domestic rabbits and hares that had died of unknown causes were also analysed. These samples were submitted to CSIRO for lagovirus testing either frozen or stored in an RNA stabilization solution containing 10 mM EDTA, 12.5 mM sodium citrate, and 2.65 M ammonium sulfate pH 5.2. Additional samples from healthy rabbits and hares were obtained from shot samples collected as part of routine vertebrate pest control program and lagovirus serological surveillance studies. Serological surveillance studies were conducted in the ACT and in Victoria. Animals were shot from a vehicle using a 0.22 calibre rifle targeting the head or chest. Sera and tissue samples (liver, duodenum, and bile) were collected post-mortem. Collection of GI.4-positive samples was described previously (Mahar *et al*., 2016). All work involving live animals (domestic and wild) was carried out according to the Australian Code for the Care and Use of Animals for Scientific Purposes with approval from the institutional animal ethics committee (CWLA-AEC #2016-02, #DOMRAB, ESAEC #13-10, ESAEC #10-12).

For detection of GI.1a-AusP/GI.2 recombinant viruses, we screened archival rabbit tissue samples and new samples that were submitted to CSIRO for lagovirus testing. These were predominantly liver samples but included two bone marrow samples, one muscle sample, and one kidney sample.

### Negative control samples

Negative control liver samples were obtained from known lagovirus-free domestic New Zealand White rabbits. Domestic rabbits were reared following the Australian Code for the Care and Use of Animals for Scientific Purposes, and all procedures were approved by the institutional animal ethics committee (CESAEC #DOMRAB). Does were not vaccinated against GI.1 and the colony is routinely monitored and confirmed to be free of GI.4 (Liu *et al*., 2012a, Liu *et al*., 2012b).

### RNA isolation

RNA was extracted from 20-30 mg of tissue or 50 μl of GI.1a-K5 stock virus using either the RNeasy mini kit (Qiagen, Chadstone Centre, VIC), the Maxwell 16 LEV simplyRNA tissue kit (Promega, Alexandria, NSW), or the Purelink viral RNA/DNA mini kit (Life Technologies, Scoresby, VIC) as per manufacturers’ instructions.

### Multiplex RT-PCR primer design

Primers were designed to specifically amplify fragments of either GI.1c, GI.1a-Aus, GI.1a-K5, or GI.2. Each primer pair was designed to produce an amplicon of a unique size that could be easily discriminated by agarose gel electrophoresis. Amplicon location, primer sequences, and nucleotide positions (based on Genbank accession KT280060) are shown in Table 1 and Figure 1. Additionally, rabbit specific primers targeting the *Oryctolagus cuniculus* 12S mitochondrial rRNA gene were included in the assay to confirm that RNA isolation was successful (Martín *et al*., 2009).

**Table 1.**
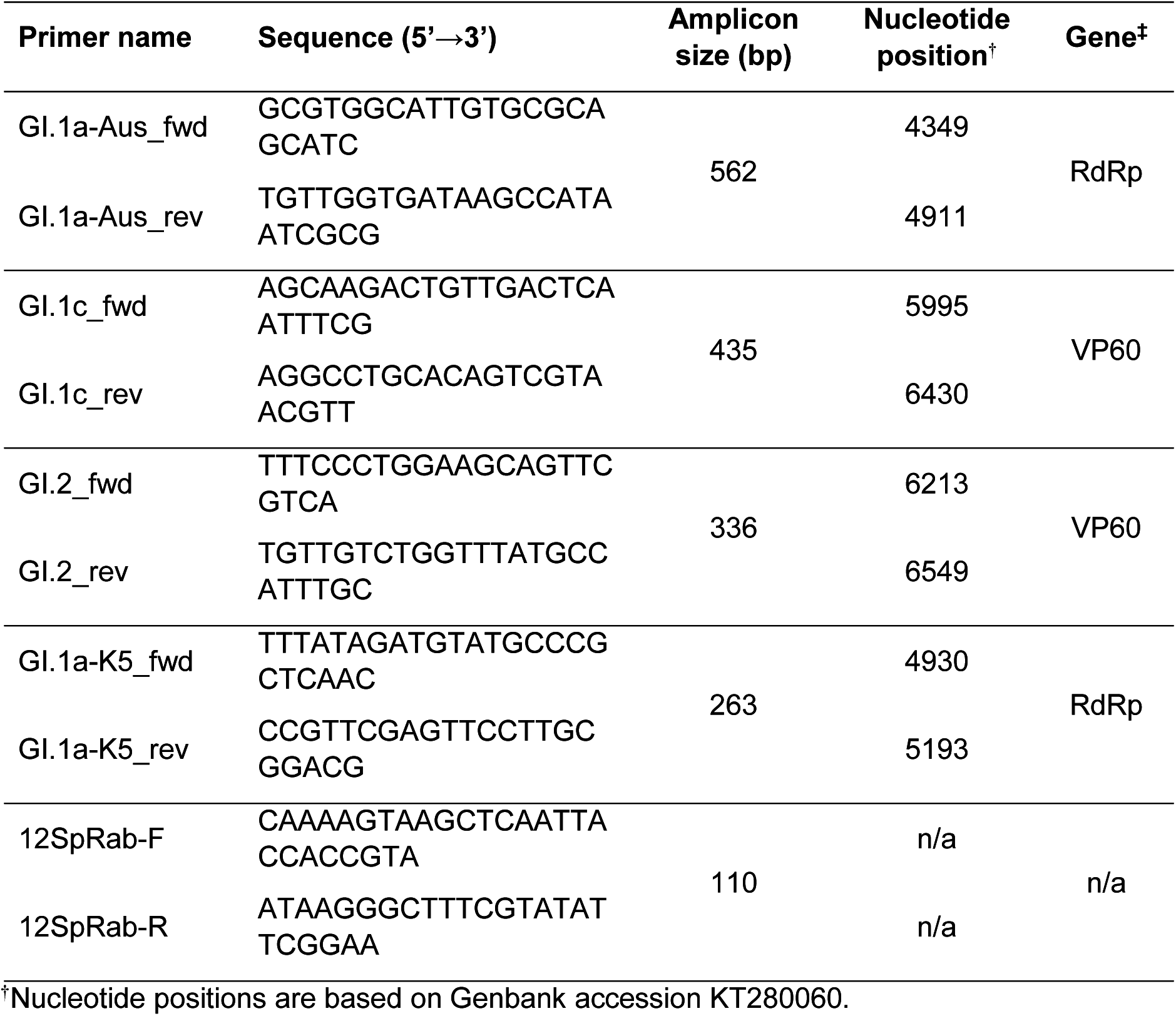
Primer sequences for the lagovirus multiplex RT-PCR assay.

### Multiplex RT-PCR development and validation

Multiplex RT-PCR was optimised using high titre RNA (6×10^7^ to 2×10^9^ copies/μl) of representative GI.1c, GI.1a-Aus, GI.1a-K5, and GI.2 viruses. All reactions were performed using the OneStep Ahead RT-PCR kit (Qiagen, Chadstone Centre, VIC). Each 10μl reaction contained 1× OneStep Ahead RT-Mix, 1× OneStep Ahead RT-PCR Master Mix, 0.5 μM each primer, and 1 μl of template RNA diluted 1/10 in nuclease free water. A ‘no template control’ and uninfected rabbit liver RNA control were included in each run. Cycling was performed using either an Eppendorf AG22331 or Applied Biosystems Proflex PCR system thermocycler, with reverse transcription being conducted at 50 °C for 10 min, followed by initial denaturation at 95 °C for 5 min, and then 40 cycles of 95 °C for 10 s, 63 °C for 20 s, 72 °C for 10 s, with a final extension at 72 °C for 2 min. The annealing temperature was optimised by gradient PCR during initial assay optimisation. Products were electrophoresed on 2% agarose gels (Bio-Rad Laboratories, Gladesville, NSW) in 1x tris-acetate EDTA, and products were visualised using SYBRsafe DNA gel stain (Life Technologies, Scoresby, VIC) on an Alpha Innotech FluorChem 8800 imaging system (Alpha Innotech, San Leandro, CA).

The specificity of the multiplex RT-PCR assay was tested using 79 known-positive tissue samples for which the virus strain had previously been determined at the EMAI veterinary virology diagnostic laboratory. The 79 samples included GI.1c (n=6), GI.1a-Aus (n=21), GI.1a-K5 (n=1), and GI.2 (n=51) viruses. Additionally, 12 domestic rabbit liver samples known to be negative for lagoviruses were screened.

Sensitivity was determined by diluting a representative GI.1c, GI.1a-Aus, and GI.2 virus to 1×10^8^ capsid copies per μl, and GI.1a-K5 to 1×10^6^ capsid copies per μl, after quantification by qRT-PCR, and preparing a 10-fold dilution series of these viruses to 1×10^1^ capsid copies per μl. Each dilution series was confirmed with qRT-PCR to ensure quantification was correct.

### qRT-PCR primer design

Primers were designed to amplify a region of VP60 that is conserved in all four pathogenic GI lagoviruses present in Australia. Primer sequences and nucleotide positions (based on Genbank accession KT280060) are shown in Table 2.

**Table 2.**
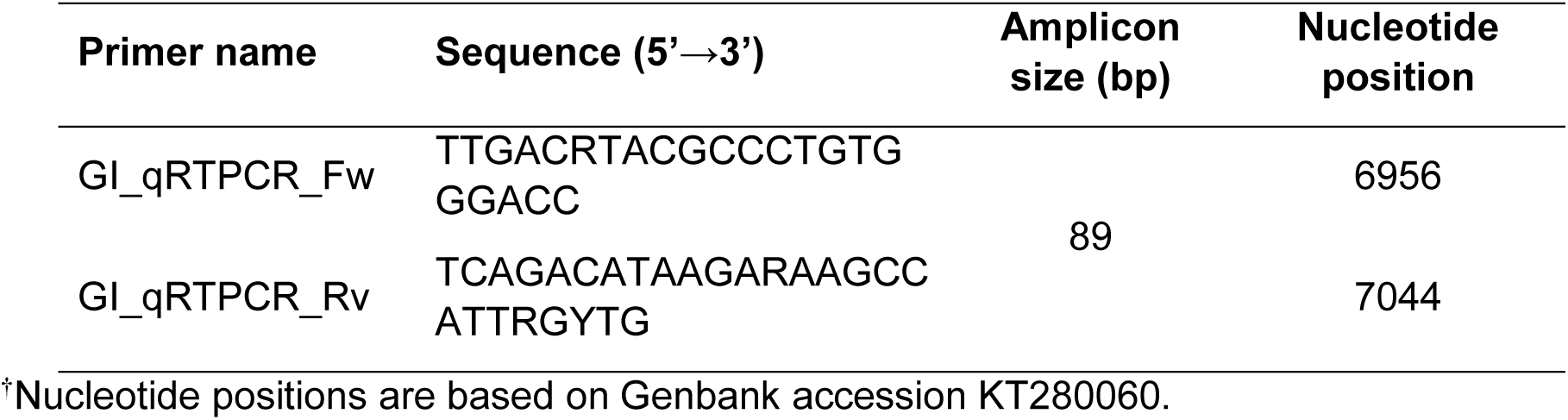
Primer sequences for the lagovirus qRT-PCR assay.

### Standards for quantification of GI lagoviruses using qRT-PCR

Full-length GI.1 genomic standards for absolute quantification of viral loads were generated by *in vitro* transcription. A plasmid construct containing the full-length GI.1c CAPM V-351 genome under control of an Sp6 promoter (Urakova *et al*., 2015) was digested with NotI-HF (Genesearch, Arundel, QLD) for 6 hours at 37°C. Digested DNA was precipitated with EDTA and sodium acetate, pH 5.5, using standard methods. DNA was quantified using the Qubit dsDNA BR assay (Life Technologies, Scoresby, VIC) and *in vitro* transcripts were prepared from 1 μg of digested DNA using the Riboprobe Combination System-SP6/T7 RNA Polymerase (Promega, Alexandria, NSW) as per manufacturer’s instructions. Transcripts were purified using the RNeasy mini kit (Qiagen, Chadstone Centre, VIC) with on-column DNase digestion, as per manufacturer’s directions. Transcripts were quantified using a NanoDrop spectrophotometer in duplicate and assessed for quality by agarose gel electrophoresis. Transcript copy number per μl of stock was determined using the following equation, where 340 g/mol was assigned as the average weight of a ribonucleotide:

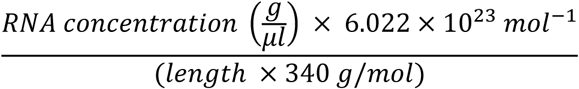

Transcript stocks were diluted to 1×10^10^ copies/μl and stored at -80 °C. Absolute quantification was performed by preparing 10-fold serial dilutions of transcripts in nuclease-free water containing 125 ng of yeast tRNA (Life Technologies, Scoresby, VIC).

### qRT-PCR optimisation

Quantitative real-time RT-PCR conditions were optimised with GI.1c *in vitro* transcripts. Reactions were performed in a final volume of 10μl and contained 1× SensiFAST SYBR No-ROX One-Step mix (Bioline, Alexandria, NSW), 0.5 μM of each primer, 0.2 μl of RNase inhibitor, 0.1 μl of reverse transcriptase, and 1 μl of template RNA. Cycling was performed using a CFX96 C1000 real-time PCR detection system (Bio-Rad Laboratories, Gladesville, NSW), with reverse transcription conducted at 45 °C for 10 min, followed by denaturation at 95 °C for 5 min, and then 40 cycles of 95 °C for 10 s, 63 °C for 40 s, 78 °C for 10 s with data acquisition. Melt curve analysis was conducted at 65-95 °C in 0.5 °C increments at 5 s per increment. Annealing temperature was optimised by gradient PCR. Data were analysed using CFX manager 3.1 software (Bio-Rad Laboratories, Gladesville, NSW) using a baseline threshold of 200 and baseline subtracted curve fit setting. Average assay efficiency was 95%. Amplicons were separated on 2% agarose gels and visualised by staining with SYBRsafe DNA gel stain (Life Technologies, Scoresby, VIC) to verify that products were of the expected size.

### qRT-PCR validation

Reactions were performed in duplicate and each run included a dilution series of full-length GI.1c transcript standards ranging from 1×10^8^ copies/μl to 1×10^2^ copies/μl for quantification, a ‘no template control’ to detect contamination, and a positive control (stored in single use aliquots) to control for inter-assay variation. Wells with melt curve peaks below 80 °C were excluded since these peaks represented primer dimer formation. Samples were excluded from analysis and repeated if the threshold cycle (Cq) standard deviation was >0.5, and runs were repeated if the positive control Cq deviated by more than 0.5 from the back-calculated average of this sample.

The qRT-PCR assay was validated initially using the 79 known lagovirus-positive tissue samples as previously determined at EMAI, including GI.1c (n=6), GI.1a-Aus (n=21), GI.1a-K5 (n=1), and GI.2 (n=51) viruses, and 12 known-negative domestic rabbit liver samples. Subsequently, additional GI.1c (n=20) and GI.2 (n=99) positive samples, as determined by the multiplex RT-PCR assay, and healthy shot rabbit or hare liver samples (n=50) were screened.

### Confirmation of a novel GI.1a-AusP/GI.2 recombinant

Sanger sequencing over the RdRp-VP60 junction was conducted to verify putative recombination in virus samples that were positive for both GI.1a-Aus and GI.2 on multiplex RT-PCR assay. First-strand cDNA was synthesised from 5 μl of RNA using 500 ng of OligodT(18mer) (GeneWorks, Thebarton, SA) and Invitrogen Superscript III or Superscript IV (Life Technologies, Scoresby, VIC) according to manufacturers’ directions. PCR was conducted using primers RHDV 11 and RHDV 12-rev (Elsworth *et al*., 2014) or GI.1a-Aus_fwd (Table 1) and GI.2-qRTPCR-R (5’-GTCAAATGTACGCTGGCTGG-3’), as described previously (Mahar *et al*., 2016). PCR amplicons were submitted to the ACRF Biomolecular Resource Facility (Canberra, ACT) for Sanger sequencing, and sequences have been deposited in Genbank under accession numbers MF598303 to MF598338. Sequences were aligned with a representative GI.1a-Aus genome (Genbank ref KY628309) and an Australian GI.2 genome (Genbank accession KT280060) using MAFFT as implemented in Geneious v9.1.6 (Kearse *et al*., 2012).

### Full genome sequencing of the novel GI.1a-AusP/GI.2 recombinant

The complete genomes of GI.1a-K5 and a suspected GI.1a-AusP/GI.2 recombinant, Car-3, were amplified in overlapping fragments and DNA libraries were prepared and sequenced using Illumina Miseq technology as described previously (Eden *et al*., 2015, Mahar *et al*., 2016, Hall *et al*., 2017). Sequences were deposited in Genbank under accession numbers MF598301 and MF598302, respectively.GI.1a-K5 was amplified using the primer pairs RHDV-1/RHDVR 6-rev; RHDV 7/RHDVR 10-rev; and RHDV 11/RHDV_end, while Car-3 was amplified using the primer pairs RHDV-1/RHDVR 6-rev; RCVf1.5/RCVr3.3; RCVf3.0/RCVr4.7; RHDV 11/RHDVR 12-rev; and RHDV f2/RHDV_end (Elsworth *et al*., 2014, Mahar *et al*., 2016, Hall *et al*., 2017). Consensus sequences were constructed by mapping cleaned reads to the lagovirus GI reference sequence (Genbank accession M67473.1 DEU/FRG/1988.50) using the Geneious package v8.1.5 (Kearse *et al*., 2012).

### Phylogenetic and recombination analyses

To explore evidence of recombination in Car-3, the complete genome sequence was aligned with 30 representative lagovirus sequences obtained from Genbank (http://www.ncbi.nlm.nih.gov/GenBank/index.html), for phylogenetic and recombination analyses. The RDP, GENECONV, MaxChai, and Bootscan methods, as available in the Recombination Detection Program v4 (Martin *et al*., 2015), were employed for recombination screening, and significant evidence of recombination was denoted by a p value <0.05. For phylogenetic analysis, a European brown hare syndrome virus (EBHSV, now referred to as GII.1) genome sequence (Genbank accession KC832839) was used as an outgroup, and the alignment was divided into non-structural genes (5,238 nt) and structural genes (2,083 nt). A maximum likelihood phylogeny was inferred for both non-structural and structural gene datasets using PhyML as available in Geneious v 8.1.5, using the GTR+I substitution model, with five rate categories, an estimated gamma distribution parameter, and a combination of nearest-neighbour interchange and subtree pruning and regrafting (SPR) branch swapping topology searching. Branch support was estimated using 1,000 bootstrap replicates and trees were rooted using the EBHSV sequence.

## Results

### Multiplex RT-PCR

A one-step multiplex RT-PCR was designed to diagnose fatal RHD and identify the causative virus, based on the pathogenic lagoviruses known to be circulating in Australia. Amplicons for GI.1c, GI.1a-Aus, GI.1a-K5, and GI.2 lagoviruses, along with a rabbit-specific amplicon to verify successful RNA extraction, could clearly be discriminated by agarose gel electrophoresis (Figure 2).

**Figure 2:**
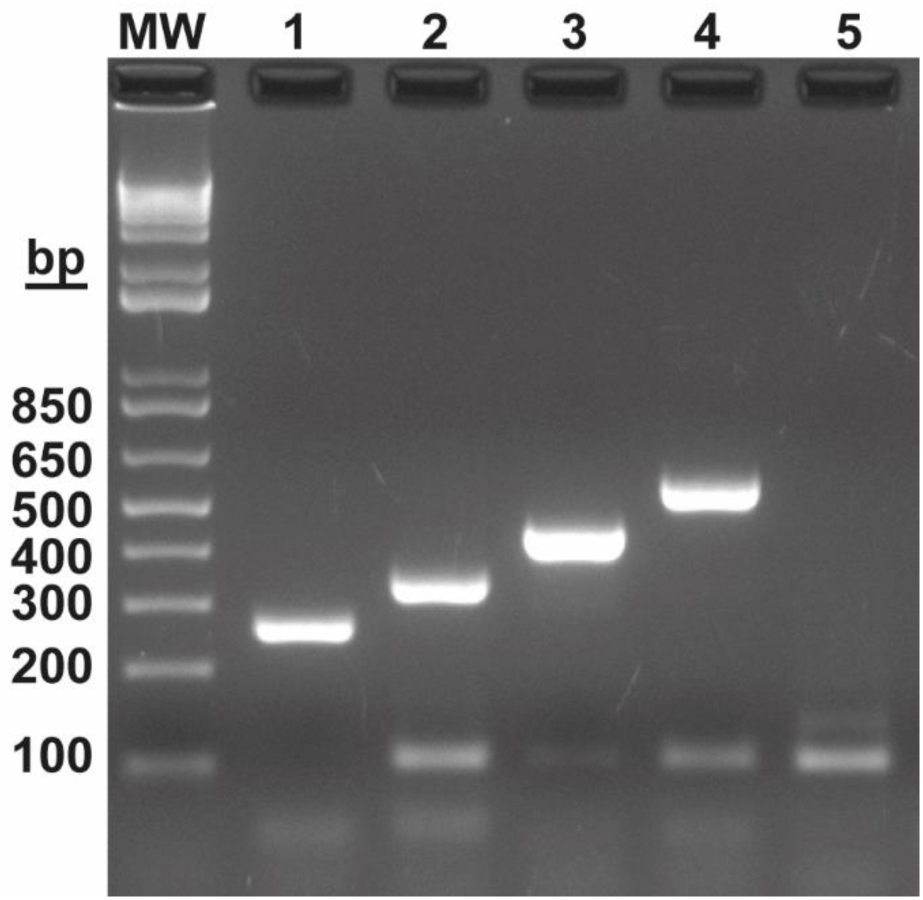
Multiplex RT-PCR for the differentiation of pathogenic GI lagoviruses in Australia. RNA was isolated from tissues of rabbits with suspected GI lagovirus infections and multiplex RT-PCR was performed to discriminate between GI.1a-K5 (1), GI.2 (2), GI.1c (3), and GI.1a-Aus (4). MW, 1 Kb Plus DNA ladder (Life Technologies, Scoresby, VIC); 5, lagovirus-negative rabbit liver; 6, no template control. A 110 bp fragment of rabbit host RNA was also amplified to confirm that RNA isolation was successful. Note that GI.1a-K5 RNA was prepared from a purified virus preparation and thus no rabbit amplicon is present.

Sensitivity of the multiplex RT-PCR assay was determined using 10-fold serial dilutions of representative samples of GI.1c, GI.1a-Aus, GI.1a-K5, and GI.2 viruses after quantification by qRT-PCR (Figure 3). The assay was able to detect all viruses down to 1×10^1^ capsid copies per μl of RNA template under these conditions.

**Figure 3:**
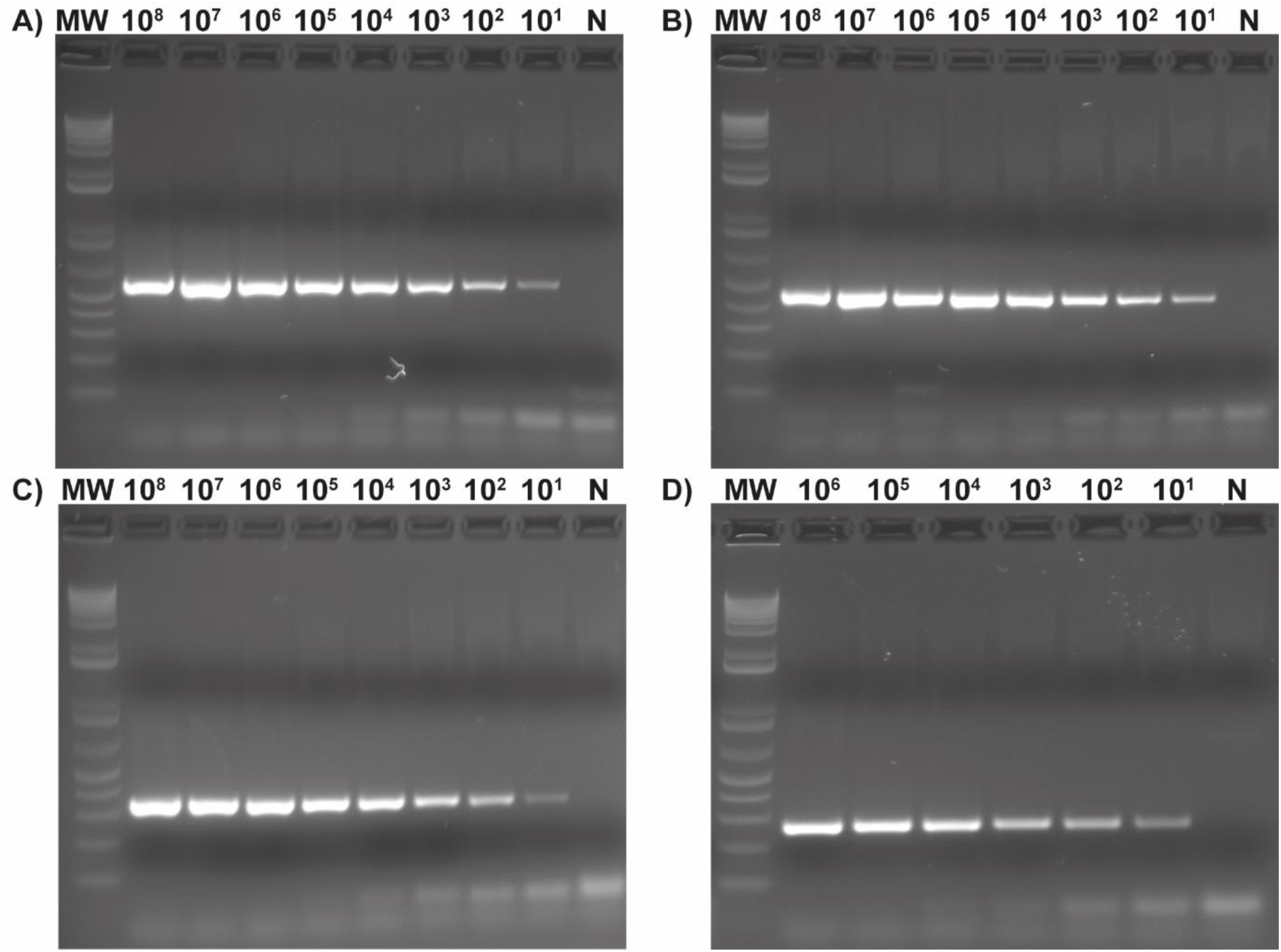
Sensitivity testing of the multiplex RT-PCR assay. Sensitivity was determined using 10-fold serial dilutions ranging from 1×10^8^ capsid copies (1×10^6^ for GI.1a-K5) to 1×10^1^ capsid copies of a representative virus of a) GI.1a-Aus; b) GI.1c; c) GI.2; and d) GI.1a-K5. MW, 1 Kb Plus DNA ladder (Life Technologies, Scoresby, VIC); N, no template control.

The multiplex RT-PCR was validated using 12 rabbit liver samples known to be lagovirus-negative, and 79 known-positive samples for which virus identification had previously been performed at an external diagnostic laboratory (EMAI). Overall, 97% (77/79) of the known-positive samples and 92% (11/12) of known-negative samples were correctly identified (Table 3). One GI.1c and one GI.2 sample were not detected by the multiplex assay, however, these samples showed very low virus titres at EMAI (Cq 34 and 33, respectively). One known-negative sample repeatedly produced a very weak amplicon approximately the size of the GI.2 product on multiplex RT-PCR assay. However, this sample was collected in 2011, before GI.2 was present in Australia, confirming that this was a non-specific amplification product. This weak non-specific product was not observed in other negative samples. Additionally, one GI.2 sample produced both a GI.2 and GI.1a-Aus band in the multiplex RT-PCR, suggesting either a mixed infection, or a possible recombinant virus.

**Table 3.** Validation of the multiplex RT-PCR assay.

| EMAI<br>genotype | Multiplex RT-PCR genotype |  |  |  |  | Total<br>No. tested |
| --- | --- | --- | --- | --- | --- | --- |
|  | GI.1c | GI.2 | GI.1a-Aus | GI.1a-K5 | Negative |  |
| GI.1c | 5 | 0 | 0 | 0 | 1 | 6 |
| GI.2 | 0 | 50<sup>‡</sup> | 0 | 0 | 1 | 51 |
| GI.1a-Aus | 0 | 0 | 21 | 0 | 0 | 21 |
| GI.1a-K5 | 0 | 0 | 0 | 1 | 0 | 1 |
| Negative<sup>†</sup> | 0 | 1<sup>§</sup> | 0 | 0 | 11 | 12 |

†Negative samples were collected from rabbits from a known lagovirus-free breeding colony.
‡One sample produced both a GI.2 and GI.1a-Aus band in multiplex RT-PCR. This was subsequently confirmed to be a novel recombinant variant.
§This sample produced a very weak non-specific amplification product approximately the size of the GI.2 amplicon despite being collected in 2011, prior to the incursion of GI.2 into Australia.
Grey shading indicates correct identification by the multiplex RT-PCR assay.

To test the ability of the multiplex RT-PCR assay to detect mixed infections, seven liver samples that were positive for both GI.1c and GI.2, as determined at EMAI, were screened. Although all samples were positive on multiplex RT-PCR, frequently only one amplicon was present, which corresponded to the dominant variant in the mixed infection based on Cq values obtained at EMAI (R. Hall, unpublished results).

### Quantitative real-time RT-PCR

A SYBR-green qRT-PCR assay was developed for quantification of virus loads in diagnostic samples. The assay amplifies an 89 bp region within the VP60 capsid gene that is conserved in all four pathogenic GI lagoviruses present in Australia. Full-length GI.1c *in vitro* transcript standards were produced to facilitate absolute quantification of virus loads. The assay was validated firstly with the 79 known lagovirus-positive RNAs and 12 known-negative RNAs used for multiplex RT-PCR validation. Of the lagovirus-positive samples, 78/79 were positive on the qRT-PCR assay, with virus loads ranging from 9×10^1^ to 4×10^9^ capsid copies per mg of tissue (x̄g = 3×10^8^). Only five samples had virus loads less than 1×10^6^ capsid copies per mg of tissue. Both the sample that was negative on our qRT-PCR and very weak positive sample (9×10^1^ capsid copies per mg of tissue) showed extremely low virus titres at EMAI (Cq 33 and 34, respectively). In all 12 of the known-negative domestic rabbit liver samples virus loads were below 10 capsid copies per mg of tissue, outside the range of the standard curve.

Subsequently, further validation was conducted with additional GI.1c (n=20) and GI.2 (n=99) positive samples, as determined by the multiplex RT-PCR assay. All 119 samples were positive on qRT-PCR. Virus loads in these samples ranged from 6×10^3^ to 8×10^9^ capsid copies per mg of tissue (X̄_g_ = 3×10^8^) and only two samples had virus loads less than 1×10^6^ capsid copies per mg of tissue. Liver samples collected from wild shot rabbits and hares (n=50) were also screened, and virus loads up to 3×10^3^ capsid copies per mg of tissue (x̄_g_ = 4×10^1^) were detected in these samples. Although these were presumed healthy shot rabbits and hares, these samples were not verified to be true negative samples, and it has previously been reported that lagovirus RNA can frequently be detected from healthy animals (Forrester *et al*., 2003). Finally, the qRT-PCR assay was also able to detect GI.4 RNA in duodenal samples, with virus loads ranging from 3×10^2^ to 3×10^5^ capsid copies per mg of duodenal tissue (X̄_g_ = 1×10^4^; R. Hall, unpublished data).

### Detection of a novel GI.1a-AusP/GI.2 recombinant

During validation of the multiplex RT-PCR, one sample, collected from NSW in July 2016 and typed as GI.2 at EMAI, produced amplicons for both GI.1a-Aus and GI.2 viruses (Figure 4). Sanger sequencing of a single amplicon spanning the RdRp-VP60 junction, the most common recombination breakpoint for caliciviruses (Bull *et al*., 2007), confirmed the presence of a recombinant virus, rather than a mixed GI.1a-Aus/GI.2 infection. Screening of archival samples and newly collected field samples using the multiplex RT-PCR assay and confirmatory Sanger sequencing detected an additional 35 cases of infection with this novel recombinant between July 2016 and June 2017, with cases occurring in NSW, the ACT, and in Victoria (Figure 5, Table 4). Interestingly, one of these recombinant detections was from a liver sample collected from a European brown hare (*Lepus europaeus*) found dead in NSW. Additionally, one case occurred in a domestic rabbit that had been vaccinated with Cylap^®^ RCD vaccine (Zoetis, West Ryde, NSW) approximately four months prior to infection.

**Figure 4:**
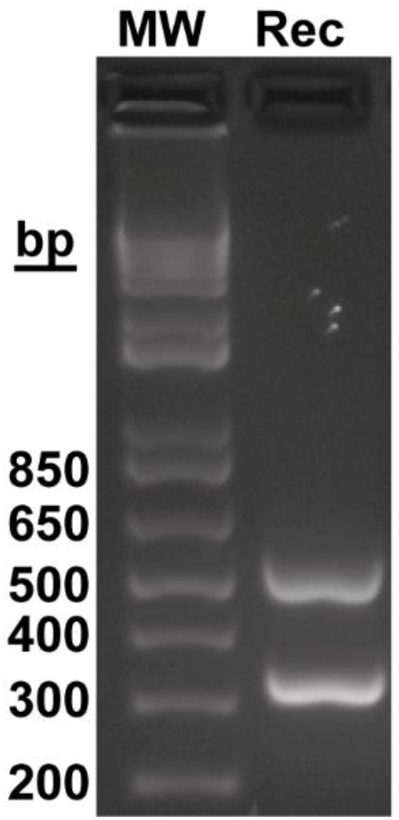
Detection of a GI.1a-AusP/GI.2 recombinant. Multiplex RT-PCR performed on RNA isolated from the liver of a rabbit suspected to have died from RHD produced amplicons for both GI.2 and GI.1a-Aus viruses, suggesting either a mixed infection or a possible recombinant virus.

**Figure 5:**
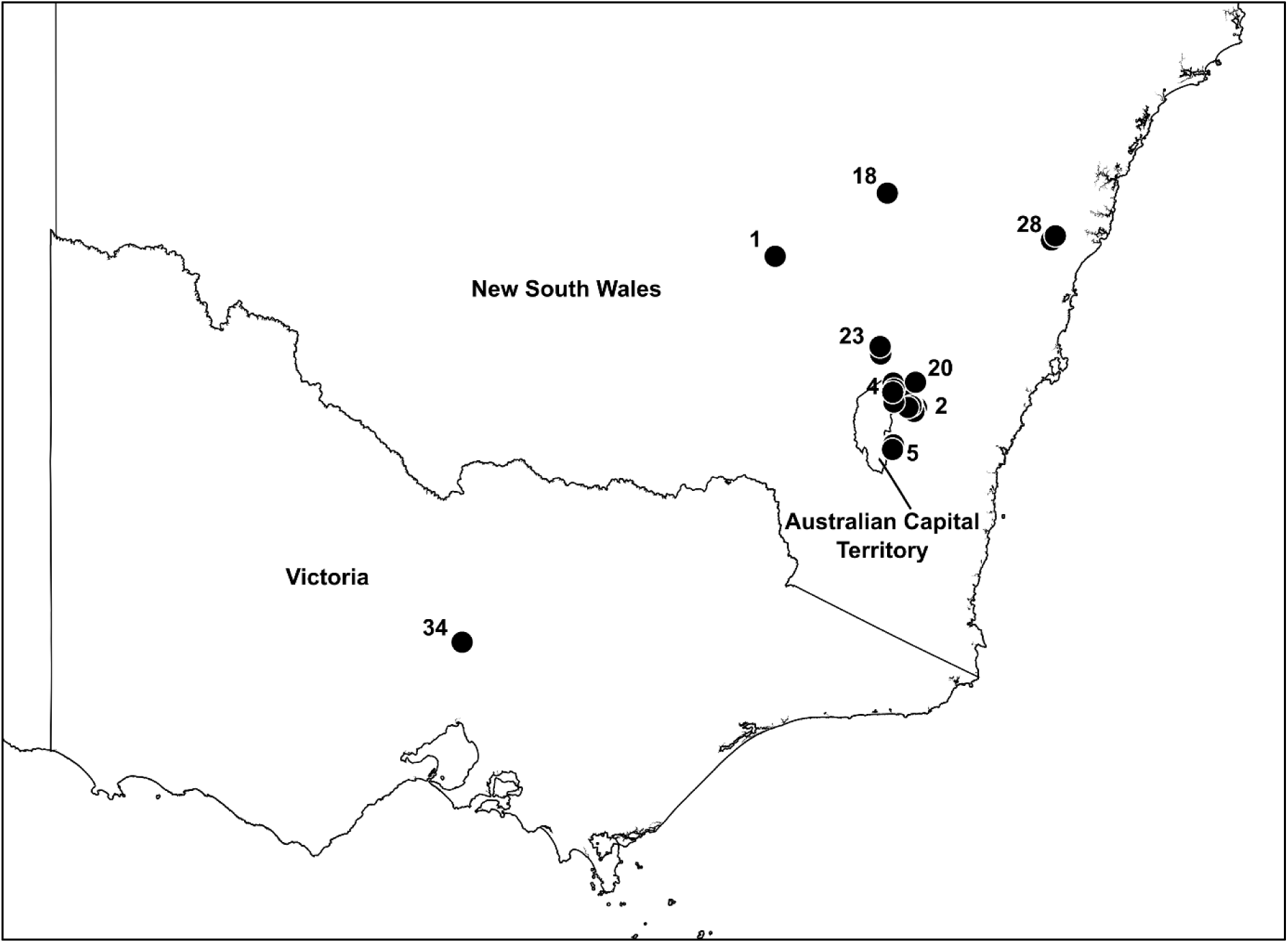
Detections of the novel GI.1a-AusP/GI.2 recombinant in Australia between July 2016 and June 2017. Sites where GI.1a-AusP/GI.2 cases were detected are indicated on the map and numbered according to the order in which the outbreaks occurred. Where multiple cases occurred in the same geographical location, only the first number is given.

**Table 4.** GI.1a-AusP/GI.2 recombinant detections, July 2016 to June 2017.

| **Case**<br>**no. <sup>†</sup>** | **Isolate name** | **Collection**<br>**date** | **Location, state<sup>‡</sup>** | **Animal**<br>**use <sup>§</sup>** |
| --- | --- | --- | --- | --- |
| 1 | AUS/NSW/TUB-1 | 13/07/2016 | Tubbul, NSW | W |
| 2 | AUS/NSW/CAR-2 | 7/10/2016 | Carwoola, NSW | W |
| 3 | AUS/NSW/CAR-3 | 10/10/2016 | Carwoola, NSW | W |
| 4 | AUS/ACT/GGH-1 | 20/11/2016 | Gungahlin, ACT | U |
| 5 | AUS/NSW/MCH-1 | 1/01/2017 | Michelago, NSW | U |
| 6 | AUS/NSW/MCH-2 | 2/01/2017 | Michelago, NSW | U |
| 7 | AUS/NSW/CAR-7 | 17/03/2017 | Carwoola, NSW | W |
| 8 | AUS/NSW/CAR-8 | 17/03/2017 | Carwoola, NSW | W |
| 9 | AUS/NSW/CAR-9 | 20/03/2017 | Carwoola, NSW | W |
| 10 | AUS/NSW/CAR-10 | 23/03/2017 | Carwoola, NSW | W |
| 11 | AUS/NSW/CAR-11 | 23/03/2017 | Carwoola, NSW | W |
| 12 | AUS/NSW/CAR-13 | 25/03/2017 | Carwoola, NSW | W |
| 13 | AUS/NSW/CAR-14 | 26/03/2017 | Carwoola, NSW | W |
| 14 | AUS/NSW/CAR-12 | 28/03/2017 | Carwoola, NSW | W |
| 15 | AUS/NSW/CAR-17 | 28/03/2017 | Carwoola, NSW | W |
| 16 | AUS/NSW/CAR-16 | 29/03/2017 | Carwoola, NSW | W |
| 17 | AUS/NSW/CAR-15 | 31/03/2017 | Carwoola, NSW | W |
| 18 | AUS/NSW/MND-1 | 10/04/2017 | Mandurama, NSW | W |
| 19 | AUS/NSW/MCH-3 | 13/04/2017 | Michelago, NSW | U |
| 20 | AUS/NSW/BYW-1 | 14/04/2017 | Bywong, NSW | U |
| 21 | AUS/ACT/HACK-3 | 15/04/2017 | Hackett, ACT | W |
| 22 | AUS/ACT/GUD-302 | 19/04/2017 | Symonston, ACT | W |
| 23 | AUS/NSW/BG-14 | 22/04/2017 | Murrumbateman, NSW | W |
| **24** | AUS/NSW/BG-15 | 25/04/2017 | Murrumbateman, NSW | W |
| **25** | AUS/NSW/BG-16 | 25/04/2017 | Murrumbateman, NSW | W |
| **26** | AUS/NSW/BG-17 | 25/04/2017 | Murrumbateman, NSW | W |
| **27** | AUS/ACT/AIN-5 | 27/04/2017 | Mt Ainslie, ACT | W |
| **28** | AUS/NSW/CAD-1 | 28/04/2017 | Camden, NSW | D |
| **29** | AUS/NSW/BG-18 | 30/04/2017 | Murrumbateman, NSW | W |
| **30** | AUS/NSW/BG-19 | 30/04/2017 | Murrumbateman, NSW | W |
| **31** | AUS/NSW/BG-20 | 30/04/2017 | Murrumbateman, NSW | W |
| **32** | AUS/NSW/BG-21 | 5/05/2017 | Murrumbateman, NSW | W |
| **33** | AUS/NSW/MCH-4 | 17/05/2017 | Michelago, NSW | W |
| **34** | AUS/VIC/KIL-1 | 27/05/2017 | Kilmore, VIC | W |
| **35** | AUS/NSW/HRP-1 | 29/05/2017 | Harrington Park, NSW | DV |
| **36** | AUS/NSW/MUR-6 | 25/06/2017 | Murrumbateman, NSW | H |

<sup>†</sup> Samples are numbered according to the date on which they were collected. These numbers correspond to those used on the map in Figure 6.
<sup>‡</sup>Abbreviations for Australian states and territories are as follows: ACT, Australian Capital Territory; NSW, New South Wales; VIC, Victoria.
<sup>§</sup>W, wild rabbit; U, animal use not specified; D, domestic rabbit; DV, domestic rabbit vaccinated with Cylap® RCD vaccine (Zoetis, West Ryde, NSW); H, hare.

Subsequently, the full genome of one of these recombinant viruses was sequenced and recombination and phylogenetic analyses were performed (Figure 6). There was significant evidence for recombination between the RdRp and VP60 genes detected by all methods in the Recombination Detection Program, with the specific breakpoint predicted to occur at nucleotide position 5,297 (equivalent to position 5,306 according to GI reference genome numbering, Genbank accession M67473.1). Phylogenetic analyses demonstrated that the non-structural genes of the novel recombinant were closely related to GI.1a-Aus sequences, sharing >98% nucleotide identity in this region, while the structural genes clustered with Australian GI.2 viruses, sharing 98.5% nucleotide identity with the structural genes of the prototype Australian GI.2, BlMt-1, with strong bootstrap support (Figure 6). Based on the newly proposed nomenclature for lagoviruses (Le Pendu *et al*., 2017), the GI.1a-Aus non-structural sequences are closely related to, but divergent from, previously described GI.4a, b, c, and d variants, thus we have designated them as GI.4e. The close relationship of the novel recombinant with Australian variants of GI.1a and GI.2, suggest that the recombination event occurred in Australia.

**Figure 6:**
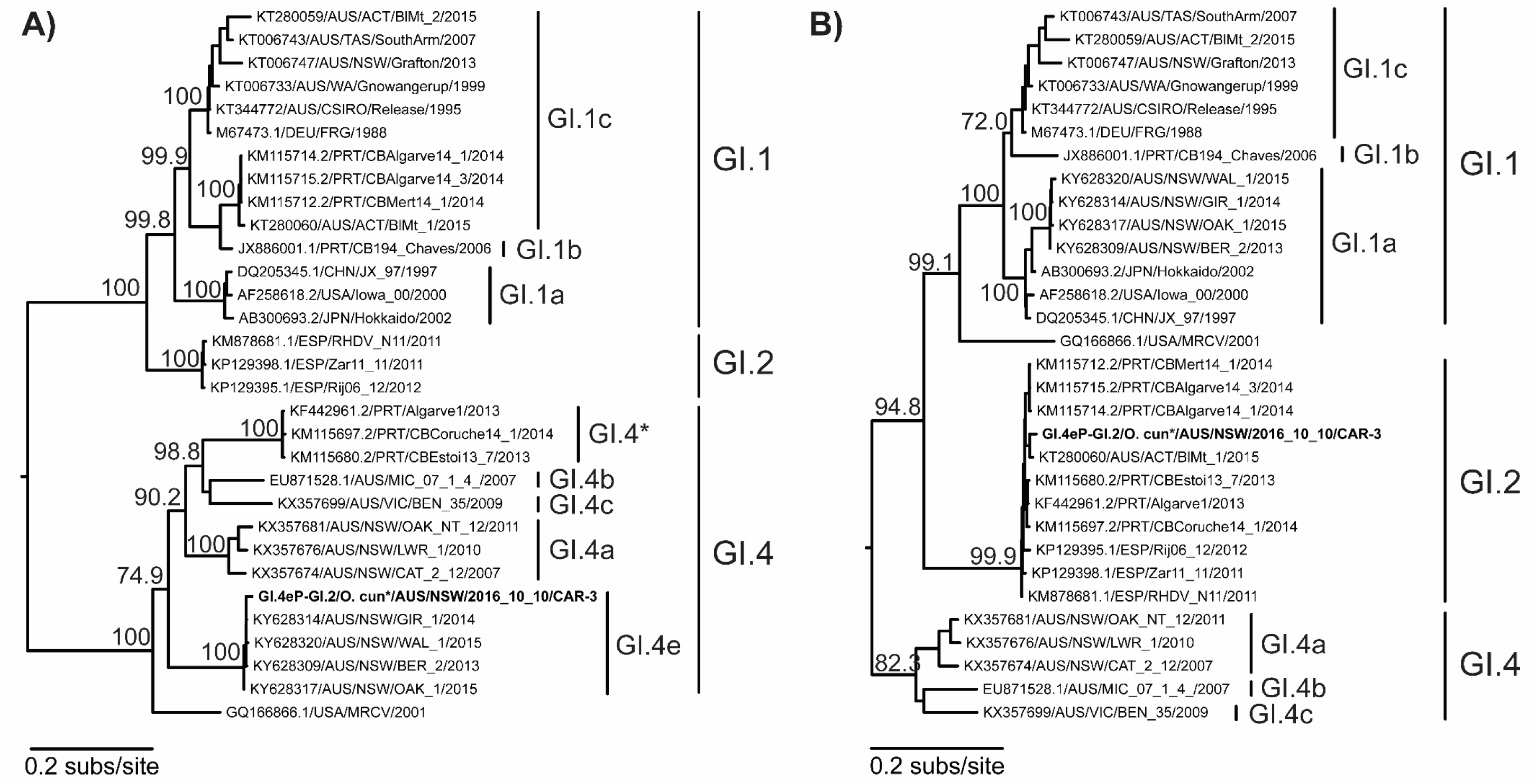
Non-structural and structural gene phylogenies of representative lagoviruses. Maximum likelihood phylogenies of the (A) nonstructural genes (n=31) and (B) structural genes (VP60 and VP10, n=31) were inferred using the newly sequenced GI.1a-AusP/GI.2 recombinant (shown in bold) and representative published sequences. The Genbank accession numbers of published sequences are indicated in the taxa names. The genotype of each cluster is indicated. The clade marked with an asterisk has not been classified to the variant level. Phylogenies were rooted using an early European EBHSV isolate (not shown) and the scale bar is proportional to the number of nucleotide substitutions per site. Bootstrap support values are shown for the major nodes.

## Discussion

We have developed a sensitive and specific multiplex RT-PCR for the detection and discrimination of pathogenic GI lagoviruses present in Australia, namely classical GI.1c, GI.2, GI.1a-Aus, and GI.1a-K5 viruses (Figure 2). In addition, we describe a SYBR-green based qRT-PCR assay for quantification of genogroup GI lagovirus loads in diagnostic samples. We have targeted the VP60 capsid gene in this assay to maximise sensitivity, since both genomic and subgenomic RNAs will be detected by these primers.

During validation of the multiplex RT-PCR assay, no viruses were misidentified as an incorrect strain, highlighting the specificity of this assay. Although a weak product was amplified from one known-negative rabbit liver sample, this was clearly distinguishable from the intense specific bands observed for known-positive samples. No amplification was detected in the negative sample by qRT-PCR, highlighting the synergism of the two assays. Mixed GI.1c/G1.2 infections were not reliably detected using the multiplex RT-PCR assay described here, since only the dominant virus was amplified consistently. However, the multiplex assay can be run in singleplex format for each virus if the index of suspicion is high for a mixed infection. It must also be noted that during routine use of the multiplex RT-PCR assay, three samples (out of over 450 positive samples tested to date) were negative on multiplex RT-PCR that were subsequently shown to have very high viral titres by qRT-PCR. When the multiplex assay was repeated, on both the initial and repeated RNA extractions, these samples returned false negative results in up to two of seven replicates. It is unknown why these templates repeatedly produced false negative results, however, all three samples were from the same RHD outbreak so sample collection, or handling and storage during transport, may have been a contributing factor. These false negatives are easily detected when the multiplex and qRT-PCR assay are used together.

The average (geometric mean) viral load in the livers of infected animals, as measured by our qRT-PCR assay, was 3 ×10^8^ capsid copies per mg of tissue. This supports previous observations that acute RHD is associated with high levels of virus replication in the liver (Elsworth *et al*., 2014). Samples with lower viral loads may represent animals that succumbed to sequelae of viral infection after a prolonged disease, or may simply be due to sample degradation, since some samples were collected from wild rabbit carcasses that were located in unfavourable environmental conditions, for example exposed to direct sunlight for an unknown period of time. Two samples that had previously been identified as weak positive samples at EMAI were either negative or had a very low virus load (9×10^1^ capsid copies per mg of tissue) outside the range of the standard curve in our qRT-PCR assay. These samples were also negative on multiplex RT-PCR. The discrepancy between our results and those of the external laboratory may be due to sample degradation during storage and transport, differences in RNA extraction methods used between the laboratories, or a higher sensitivity of the Taqman assay used at the external laboratory. However, the diagnostic significance of these very low titres is questionable given that animals with terminal RHD invariably have very high viral loads (Elsworth *et al*., 2014).

Interestingly, virus loads up to 3×10^3^ capsid copies per mg of tissue were detected in the livers of presumed negative healthy wild rabbits and hares, which were negative on multiplex RT-PCR and negative for GI.4 viruses by PCR. In contrast, in true negative domestic animals that had never been exposed to known caliciviruses, fewer than 10 copies of viral RNA per mg liver tissue were detected, outside the range of the standard curve for the assay. For precise quantification of the respective strains, virus-specific Taqman qRT-PCR assays with virus-specific standards would be preferable (such as those reported previously (Gall *et al*., 2007)). However, the assay described here is designed to be used in conjunction with the highly sensitive multiplex RT-PCR assay to ascertain whether GI lagoviruses were the likely cause of death in the animals tested, which is suggested by high viral loads. In this context, the low virus titres detected in wild lagomorphs, in combination with the lack of amplification in the multiplex RT-PCR assay, may indicate the presence of one or more as-yet uncharacterised benign lagoviruses present in Australian lagomorph populations, since the qRT-PCR assay was designed to be broadly reactive. Alternatively, the qRT-PCR assay may be detecting lagovirus RNA from animals recovering from a previous non-fatal lagovirus infection, since non-infectious GI.1 RNA has been shown to be detectable for at least 15 weeks post-infection (Gall *et al*., 2007). Previous studies from New Zealand have also demonstrated the detection of GI.1 RNA by nested RT-PCR in a large proportion of healthy shot wild rabbits (Zheng *et al*., 2002, Forrester *et al*., 2003). In the latter case, we would also expect a positive result by multiplex RT-PCR, since the sensitivity of that assay was determined to be 1×10^1^ capsid copies per μl of RNA template for each virus strain (Figure 3). However, sensitivity testing was performed using serial dilutions of a high titre sample and the proportion of viral RNA to host RNA would be lower in a low titre sample compared to a diluted high titre sample, which may also affect the assay sensitivity.

It is notable that our qRT-PCR assay was also able to detect GI.4 benign virus RNA in duodenal samples, demonstrating the utility of this assay to detect genetically diverse GI lagoviruses. Virus loads during GI.4 infection (x̄_g_ = 1×10^4^ capsid copies per mg of tissue) were considerably lower than those detected for infection with pathogenic lagoviruses, as reported previously (Strive *et al*., 2010).

Alternative diagnostic methods, such as high-throughput sequencing technologies, have also been developed and utilised for analyses of lagoviruses in the Australian context (Eden *et al*., 2015, Hall *et al*., 2015, Mahar *et al*., 2016). However, despite recent advances these technologies remain complex, with regards to both sample preparation and data analysis, and are more expensive and slower than conventional diagnostic testing. Furthermore, the risk of contamination during preparation of sequencing libraries is extremely high, the diagnostic significance of low numbers of reads is questionable, and mixed infections are not reliably detected (R. Hall and J. Mahar, unpublished data). The assays described here, despite their limitations, constitute a fast, robust, and cost-effective diagnostic pipeline for routine rabbit GI lagovirus testing, where fast turnaround of results is required.

Using the newly developed multiplex RT-PCR assay and qRT-PCR assay, we detected 36 cases of infection with a novel recombinant virus, which had the non-structural genes of GI.1a-Aus and the structural genes of GI.2. These cases were detected in NSW, the ACT, and Victoria between July 2016 and June 2017 (Table 4, Figure 5) and confirmed by Sanger sequencing over the RdRp-VP60 junction. The detection of this recombinant was only possible because the primer binding sites for GI.1a-Aus lie within the RdRp gene while those for GI.2 lie within the VP60 region. Accordingly, the multiplex assay reported here will not be able to detect GI.1c/GI.2 recombinants or GI.1a-Aus/GI.1a-K5 recombinants, due to the relative location of the primer binding sites for these assays (Figure 1). High throughput sequencing methods spanning the RdRp-VP60 junction region may facilitate rapid detection of intergenic recombination between any rabbit lagoviruses.

Although sampling of dead rabbits is not systematic, it seems clear that this recombinant has already spread further than the parental GI.1a-Aus virus. It is likely that this recombinant virus will have similar immunological characteristics to GI.2, however, experimental determination of relative virulence and transmissibility have not yet been determined. This immunological similarity is supported by the detection of this recombinant virus in a domestic rabbit that had been vaccinated against GI.1 strains approximately four months prior to infection. Interestingly, this recombinant variant was also detected in a hare, suggesting that the GI.2 capsid protein is responsible for permissivity of hare cells to infection. This once again highlights the importance of recombination for generating genetic diversity in caliciviruses and indicates the frequency with which new viruses can emerge when genetically diverse lagoviruses circulate concurrently. The discovery of this new recombinant virus emphasises the importance of maintaining monitoring efforts to remain informed about the way lagoviruses are evolving and interacting within Australia.

We report an improved diagnostic testing method for rapid detection and discrimination of pathogenic rabbit lagoviruses in the Australian context. The multiplex RT-PCR described here is a sensitive and specific assay that can rapidly diagnose infection with pathogenic lagoviruses and discriminate between GI.1c, GI.2, GI.1a-Aus, and GI.1a-K5 viruses. Additionally, we report a SYBR-green based qRT-PCR assay that broadly detects all of the known rabbit GI lagoviruses present in Australia and permits quantification of virus load in tissue samples. The development of these assays facilitates rapid detection, identification, and quantification of these viruses in a high throughput and cost-effective method. Since both assays are one-step RT-PCRs, there is no requirement for a separate cDNA synthesis step, reducing sample processing time and cost. These assays represent useful tools for ongoing large-scale nationwide monitoring of the spread and effectiveness of the GI.1a-K5 virus after its release in 2017, which will assist in guiding wild rabbit biocontrol programs into the future.

## Acknowledgements

We thank submitters and the Rabbitscan monitoring network for providing the samples utilised in this study. The assistance of technical staff of the EMAI virology laboratory is greatly appreciated. Funding for this work was obtained from CSIRO and the Invasive Animals Cooperative Research Centre. J.E.M. is supported by grant DP140103362 from the Australian Research Council.

## Conflict of Interest Statement

The authors declare that they have no competing interests.

## References

Abrantes, J., W. van der Loo, J. Le Pendu and P. J. Esteves, 2012: Rabbit haemorrhagic disease (RHD) and rabbit haemorrhagic disease virus (RHDV): a review. Veterinary Research, 43, 12. doi: 10.1186/1297-9716-43-12

Bull, R. A., M. M. Tanaka and P. A. White, 2007: Norovirus recombination. Journal of General Virology, 88, 3347–3359. doi: 10.1099/vir.0.83321-0

Capucci, L., F. Fallacara, S. Grazioli, A. Lavazza, M. L. Pacciarini and E. Brocchi, 1998: A further step in the evolution of rabbit hemorrhagic disease virus: the appearance of the first consistent antigenic variant. Virus Research, 58, 115–126. doi:10.1016/S0168-1702(98)00106-3

Capucci, L., P. Fusi, A. Lavazza, M. L. Pacciarini and C. Rossi, 1996: Detection and preliminary characterization of a new rabbit calicivirus related to rabbit hemorrhagic disease virus but nonpathogenic. Journal of Virology, 70, 8614–8623. doi: not available

Cooke, B. D. and F. Fenner, 2002: Rabbit haemorrhagic disease and the biological control of wild rabbits, *Oryctolagus cuniculus*, in Australia and New Zealand. Wildlife Research, 29, 689–706. doi: 10.1071/WR02010

Cox, T. E., T. Strive, G. Mutze, P. West and G. Saunders, 2013: Benefits of rabbit biocontrol in Australia. PestSmart Toolkit publication, Invasive Animals Cooperative Research Centre, Canberra, Australia.

Dalton, K. P., I. Nicieza, A. Balseiro, M. A. Muguerza, J. M. Rosell, R. Casais, Á. L. Álvarez and F. Parra, 2012: Variant rabbit hemorrhagic disease virus in young rabbits, Spain. Emerging Infectious Diseases, 18, 2009–2012. doi: 10.3201/eid1812.120341

Eden, J.-S., J. Kovaliski, J. A. Duckworth, G. Swain, J. E. Mahar, T. Strive and E. C. Holmes, 2015: Comparative phylodynamics of rabbit haemorrhagic disease virus (RHDV) in Australia and New Zealand. Journal of Virology. doi: 10.1128/JVI.01100-15

Elsworth, P., B. D. Cooke, J. Kovaliski, R. Sinclair, E. C. Holmes and T. Strive, 2014: Increased virulence of rabbit haemorrhagic disease virus associated with genetic resistance in wild Australian rabbits (*Oryctolagus cuniculus*). Virology, 464–465, 415-423. doi: 10.1016/j.virol.2014.06.037

Forrester, N. L., B. Boag, S. R. Moss, S. L. Turner, R. C. Trout, P. J. White, P. J. Hudson and E. A. Gould, 2003: Long-term survival of New Zealand rabbit haemorrhagic disease virus RNA in wild rabbits, revealed by RT-PCR and phylogenetic analysis. Journal of General Virology, 84, 3079–3086. doi: 10.1099/vir.0.19213-0

Forrester, N. L., S. R. Moss, S. L. Turner, H. Schirrmeier and E. A. Gould, 2008: Recombination in rabbit haemorrhagic disease virus: possible impact on evolution and epidemiology. Virology, 376, 390–396. doi: 10.1016/j.virol.2008.03.023

Forrester, N. L., R. C. Trout and E. A. Gould, 2007: Benign circulation of rabbit haemorrhagic disease virus on Lambay Island, Eire. Virology, 358, 18–22. doi: 10.1016/j.virol.2006.09.011

Gall, A., B. Hoffmann, J. P. Teifke, B. Lange and H. Schirrmeier, 2007: Persistence of viral RNA in rabbits which overcome an experimental RHDV infection detected by a highly sensitive multiplex real-time RT-PCR. Veterinary Microbiology, 120, 17–32. doi: 10.1016/j.vetmic.2006.10.006

Hall, R. N., L. Capucci, M. Matthaei, S. Esposito, P. J. Kerr, M. Frese and T. Strive, 2017: An *in vivo* system for directed experimental evolution of rabbit haemorrhagic disease virus. PLOS ONE, 12, e0173727. doi: 10.1371/journal.pone.0173727

Hall, R. N., J. E. Mahar, S. Haboury, V. Stevens, E. C. Holmes and T. Strive, 2015: Emerging rabbit hemorrhagic disease virus 2 (RHDVb), Australia. Emerging Infectious Diseases, 21, 2276–2278. doi: 10.3201/eid2112.151210

Hoehn, M., P. J. Kerr and T. Strive, 2013: *In situ* hybridisation assay for localisation of rabbit calicivirus Australia-1 (RCV-A1) in European rabbit (*Oryctolagus cuniculus*) tissues. Journal of Virological Methods, 188, 148–152. doi: 10.1016/j.jviromet.2012.11.043

Kearse, M., R. Moir, A. Wilson, S. Stones-Havas, M. Cheung, S. Sturrock, S. Buxton, A. Cooper, S. Markowitz, C. Duran, T. Thierer, B. Ashton, P. Meintjes and A. Drummond, 2012: Geneious Basic: an integrated and extendable desktop software platform for the organization and analysis of sequence data. Bioinformatics, 28, 1647–1649. doi: 10.1093/bioinformatics/bts199

Kovaliski, J., R. Sinclair, G. Mutze, D. Peacock, T. Strive, J. Abrantes, P. J. Esteves and E. C. Holmes, 2014: Molecular epidemiology of rabbit haemorrhagic disease virus in Australia: when one became many. Molecular Ecology, 23, 408–420. doi: 10.1111/mec.12596

Lavazza, A. and L. Capucci, 2016: Rabbit haemorrhagic disease. OIE Manual of Diagnostic Tests and Vaccines for Terrestrial Animals. http://www.oie.int/fileadmin/Home/eng/Healthstandards/tahm/2.06.02RHD.pdf.

Le Gall-Recule, G., A. Lavazza, S. Marchandeau, S. Bertagnoli, F. Zwingelstein, P. Cavadini, N. Martinelli, G. Lombardi, J. L. Guerin, E. Lemaitre, A. Decors, S. Boucher, B. Le Normand and L. Capucci, 2013: Emergence of a new lagovirus related to Rabbit haemorrhagic disease virus. Veterinary research, 44, 81. doi: 10.1186/1297-9716-44-81

Le Gall-Recule, G., F. Zwingelstein, S. Boucher, B. Le Normand, G. Plassiart, Y. Portejoie, A. Decors, S. Bertagnoli, J. L. Guerin and S. Marchandeau, 2011a: Detection of a new variant of rabbit haemorrhagic disease virus in France. Veterinary record, 168, 137–138. doi: 10.1136/vr.d697

Le Gall-Recule, G., F. Zwingelstein, M. P. Fages, S. Bertagnoli, J. Gelfi, J. Aubineau, A. Roobrouck, G. Botti, A. Lavazza and S. Marchandeau, 2011b: Characterisation of a non-pathogenic and non-protective infectious rabbit lagovirus related to RHDV. Virology, 410, 395–402. doi: 10.1016/j.virol.2010.12.001

Le Gall-Recule, G., F. Zwingelstein, S. Laurent, C. de Boisseson, Y. Portejoie and D. Rasschaert, 2003: Phylogenetic analysis of rabbit haemorrhagic disease virus in France between 1993 and 2000, and the characterisation of RHDV antigenic variants. Archives of Virology, 148, 65–81. doi: 10.1016/j.virol.2005.10.006

Le Pendu, J., J. Abrantes, S. Bertagnoli, J. S. Guitton, G. Le Gall-Recule, A. M. Lopes, S. Marchandeau, F. Alda, T. Almeida, A. P. Célio, J. Bárcena, G. Burkmakina, E. Blanco, C. Calvete, P. Cavadini, B. D. Cooke, K. P. Dalton, M. D. Mateos, W. Deptula, J. S. Eden, F. Wang, C. C. Ferreira, P. Ferreira, P. Foronda, D. Gonçalves, D. Gavier-Widen, R. N. Hall, B. Hukowska-Szematowicz, P. J. Kerr, J. Kovaliski, A. Lavazza, J. E. Mahar, A. Malogolovkin, R. M. Marques, S. Marques, A. Martin-Alonso, P. Monterroso, G. Mutze, A. Neimanis, P. Niedzwiedzka-Rystwej, D. Peacock, F. Parra, M. Rocchi, C. Rouco, N. Ruvoën-Clouet, E. Silva, D. Silvério, T. Strive, G. Thompson, B. Tokarz-Deptula, P. J. Esteves and S. Moreno, 2017: Proposal for a unified classification system and nomenclature of lagoviruses. Journal of General Virology, in press. doi:

Liu, J., P. J. Kerr and T. Strive, 2012a: A sensitive and specific blocking ELISA for the detection of rabbit calicivirus RCV-A1 antibodies. Virology Journal, 9, 182. doi: 10.1186/1743-422X-9-182

Liu, J., P. J. Kerr, J. D. Wright and T. Strive, 2012b: Serological assays to discriminate rabbit haemorrhagic disease virus from Australian non-pathogenic rabbit calicivirus. Veterinary Microbiology, 157, 345–354. doi: 10.1016/j.vetmic.2012.01.018

Liu, S. J., H. P. Xue, B. Q. Pu and N. H. Qian, 1984: A new viral disease in rabbits. Journal of Veterinary Diagnostic Investigations, 16, 253–255. doi: not available

Lopes, A. M., K. P. Dalton, M. J. Magalhaes, F. Parra, P. J. Esteves, E. C. Holmes and J. Abrantes, 2015: Full genomic analysis of new variant rabbit hemorrhagic disease virus (RHDVb) revealed multiple recombination events. Journal of General Virology, 96, 1309–1319. doi: 10.1099/vir.0.000070

Mahar, J. E., L. Nicholson, J. S. Eden, S. Duchene, P. J. Kerr, J. Duckworth, V. K. Ward, E. C. Holmes and T. Strive, 2016: Benign rabbit caliciviruses exhibit evolutionary dynamics similar to those of their virulent relatives. Journal of Virology, 90, 9317–9329. doi: 10.1128/JVI.01212-16

Martin, D. P., B. Murrell, M. Golden, A. Khoosal and B. Muhire, 2015: RDP4: Detection and analysis of recombination patterns in virus genomes. Virus Evolution. doi: 10.1093/ve/vev003

Martín, I., T. García, V. Fajardo, M. Rojas, N. Pegels, P. E. Hernández, I. González and R. Martín 2009: Polymerase chain reaction detection of rabbit DNA in food and animal feed. World Rabbit Science, 17, 27–34. doi: 10.4995/wrs.2009.667

Meyers, G., C. Wirblich and H. J. Thiel, 1991: Genomic and subgenomic RNAs of rabbit hemorrhagic disease virus are both protein-linked and packaged into particles. Virology, 184, 677–686. doi: 10.1016/0042-6822(91)90437-G

Oem, J. K., K. N. Lee, I. S. Roh, K. K. Lee, S. H. Kim, H. R. Kim, C. K. Park and Y. S. Joo, 2009: Identification and characterization of rabbit hemorrhagic disease virus genetic variants isolated in Korea. Journal of Veterinary Medical Science, 71, 1519–1523. doi: 10.1292/jvms.001519

Ohlinger, V. F., B. Haas, G. Meyers, F. Weiland and H. J. Thiel, 1990: Identification and characterization of the virus causing rabbit hemorrhagic disease. Journal of Virology, 64, 3331–3336. doi: not available

OIE, 2014: Rabbit haemorrhagic disease, Australia - immediate notification report Ref OIE = 14719.

OIE, 2017: Rabbit haemorrhagic disease, Australia - immediate notification report Ref OIE = 23377.

Schirrmeier, H., I. Reimann, B. Kollner and H. Granzow, 1999: Pathogenic, antigenic and molecular properties of rabbit haemorrhagic disease virus (RHDV) isolated from vaccinated rabbits: detection and characterization of antigenic variants. Archives of Virology, 144, 719–735. doi: not available

Strive, T., J. Wright, J. Kovaliski, G. Botti and L. Capucci, 2010: The non-pathogenic Australian lagovirus RCV-A1 causes a prolonged infection and elicits partial cross-protection to rabbit haemorrhagic disease virus. Virology, 398, 125–134. doi: 10.1016/j.virol.2009.11.045

Strive, T., J. D. Wright and A. J. Robinson, 2009: Identification and partial characterisation of a new Lagovirus in Australian wild rabbits. Virology, 384, 97–105. doi: 10.1016/j.virol.2008.11.004

Urakova, N., M. Frese, R. N. Hall, J. Liu, M. Matthaei and T. Strive, 2015: Expression and partial characterisation of rabbit haemorrhagic disease virus non-structural proteins. Virology, 484, 69–79. doi: 10.1016/j.virol.2015.05.004

Zheng, T., A. M. Napier, J. P. Parkes, J. S. O’Keefe and P. H. Atkinson, 2002: Detection of RNA of rabbit haemorrhagic disease virus from New Zealand wild rabbits. Wildlife Research, 29, 683–688. doi: 10.1071/WR01071

